# Re-evaluating the nuclear staining on pathological sections in Alzheimer’s disease

**DOI:** 10.1101/2023.03.22.533761

**Authors:** Hualin Fu, Jilong Li, Chunlei Zhang, Peng Du, Guo Gao, Qiqi Ge, Xinping Guan, Daxiang Cui

## Abstract

A paper recently published on Nature Neuroscience, asserting that lysosome leakage induces neuronal cell death and the formation of senile plaques in Alzheimer’s disease. The study believed that flower-like perikaryal rosettes formed by packed autophagic vacuoles around central nuclei, termed “PANTHOS” neurons, are responsible for the senile plaque formation. However, all of the nuclei staining in this paper based on images in the blue fluorescence channel. We did detailed studies on the senile plaque blue autofluorescence and found that senile plaque blue autofluorescence could be easily mistaken as nuclear staining. The blue “nuclear” staining in this paper is probably inaccurate since it was not verified in red or green fluorescence channels and their “nuclei”-related data might be mis-interpreted.

## Main Text

A paper recently published on Nature Neuroscience^1^, stating that lysosome leakage induced neuronal cell death and the formation of senile plaques in Alzheimer’s disease. The study believed that flower-like perikaryal rosettes formed by packed autophagic vacuoles around central nuclei, termed “PANTHOS”, are responsible for senile plaque formation. This study is a very influential study and it has been cited more than 60 times as shown from the PubMed database since its publication in June, 2022. The neural cell nuclear staining data is an important part of this research. However, we noticed that all of the nuclei staining in this paper was based on fluorescent images in the blue fluorescence channel. We did extensive studies on the senile plaque blue autofluorescence material, which was assigned a temporary name of MetaBlue in our pervious study^2^, and found that senile plaque MetaBlue autofluorescence could be easily mistaken as blue nuclei staining. Thus, the blue “nuclei” staining in this paper is probably inaccurate since it was not verified in other fluorescent channels and their “nuclei”-related data might be mis-interpreted.

Several previous studies had found that there is characteristic blue auto fluorescence in the senile plaques and also in CAA^2–6^. All of the cores of dense-core senile plaques have strong blue auto fluorescence with no exception according to our observations.

Since the blue nuclear staining cannot be differentiated from the MetaBlue auto fluorescence, accurate identification of cell nucleus is impossible from the blue fluorescence channel alone on AD brain pathological sections. In this paper^1^, DAPI blue fluorescence nuclear staining in Figure 5f, Figure 6b, Figure 6c, Figure 7c, Extended Data Figure 3a, Extended Data Figure 4, Extended Figure 9e should be double-checked with nuclear dye in red or green fluorescence channels in order to make precise interpretation of the images. In the study, they also used blue fluorescence channel for other nuclear markers such as histone H3 and Lamin A/C (Figure 4b). For the same reasons, all these blue fluorescence staining results could be confounded by the blue auto fluorescence of senile plaques. These staining should also be confirmed with green or red fluorescence secondary antibodies in order to be scientifically accurate.

We used the red fluorescence nuclei dye PI to show that the senile plaque MetaBlue auto blue fluorescence never co-localized with any nuclei staining, and vice versa, PI-labeled nuclei did not co-localize with senile plaque MetaBlue fluorescence (Figure 1). In Figure 1A, we did see some nuclear staining in the central region of a plaque, but this particular plaque center was not having central Aβ staining or central MetaBlue fluorescence. For all other plaques with core Aβ and core MetaBlue fluorescence in Figure 1, there is no PI staining in plaque cores. We did thorough investigation on the nuclei labeling of the cores of dense-core plaques on AD frontal brain sections. PI labeling of 59 dense-core plaques were recorded. None of them showed PI labeling in the plaque cores, which means that, most of the time the cores of dense-core plaques have no nuclei staining^2^. In our experiments, nuclear staining and plaque core fibrillar Aβ or MetaBlue auto fluorescence staining is mutually exclusive. When looking closely at the morphology of nuclear staining comparing to plaque core blue auto fluorescence, we could see a difference that the nuclear staining images usually have smooth boundaries whereas the amyloid core blue auto fluorescence images often have ruffled edges, likely due to the nature of amyloid plaque formation. Some of the blue images annotated as PANTHOS neuron “nuclei” had the ruffled edge appearance of MetaBlue fluorescence. In addition, in Figure 5f, we could see 10 senile plaques displaying characteristic MetaBlue fluorescence. In the article, Figure 5f, Figure 6b, Figure 6c, Figure 7c, Extended Data Figure 3a, Extended Data Figure 4, Extended Figure 9e all indicated the amyloid core was colocalizing with “PANTHOS” neuron nuclei staining, while the data in Figure 8a, 8d, Extended Data Figure 7c showed that the amyloid core was made up of amyloid materials as labeled by Thio-S staining. In another word, in this article, both nuclei staining and Thio-S-positive amyloid materials localized to the amyloid cores, which was not what we saw in our study. We did observe that some nucleated neural cells have diffusive spotted Aβ signals representing neuronal cell Aβ expression. However, neural cell Aβ staining was quite different from the aggregated amyloid plaque Aβ staining and was significantly weaker, as we analyzed in the published study^2^. We also did Cathepsin D immunostaining experiments with no nuclear counter stains. The results showed similar rosette-shape staining patterns with plaque blue auto fluorescence in the cores with Cathepsin D staining wrapped the central blue fluorescence while senile plaque Aβ mostly co-localized with the MetaBlue fluorescence (Figure 2). The central MetaBlue fluorescence in the cores of dense-core plaques can be easily mistaken as nuclei staining when using blue fluorescent nuclei dyes such as Hoechst or DAPI. In our published article, we also showed clearly that senile plaque MetaBlue fluorescence was independent of nuclear dye staining^2^.

**Figure 1:**
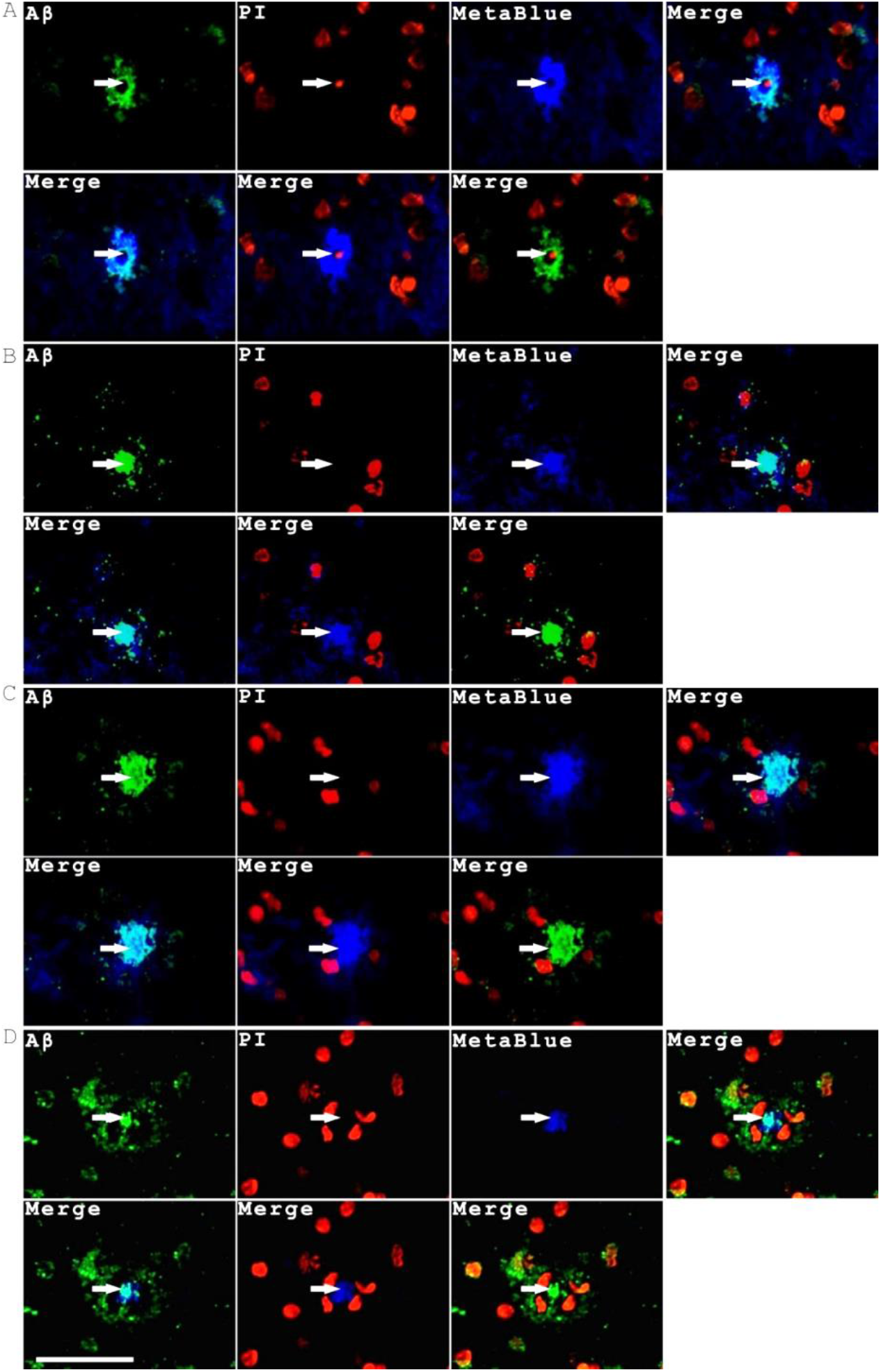
Senile plaque auto blue fluorescence (MetaBlue) is not coming from nuclear materials. MetaBlue blue auto fluorescence does not co-localize with any nucleus staining by PI in all the senile plaques shown (**A-D**). On the other hand, PI-stained nuclei do not have MetaBlue fluorescence either. The arrows indicated materials with MetaBlue blue auto fluorescence. Scale bar, 50 μm.

**Figure 2:**
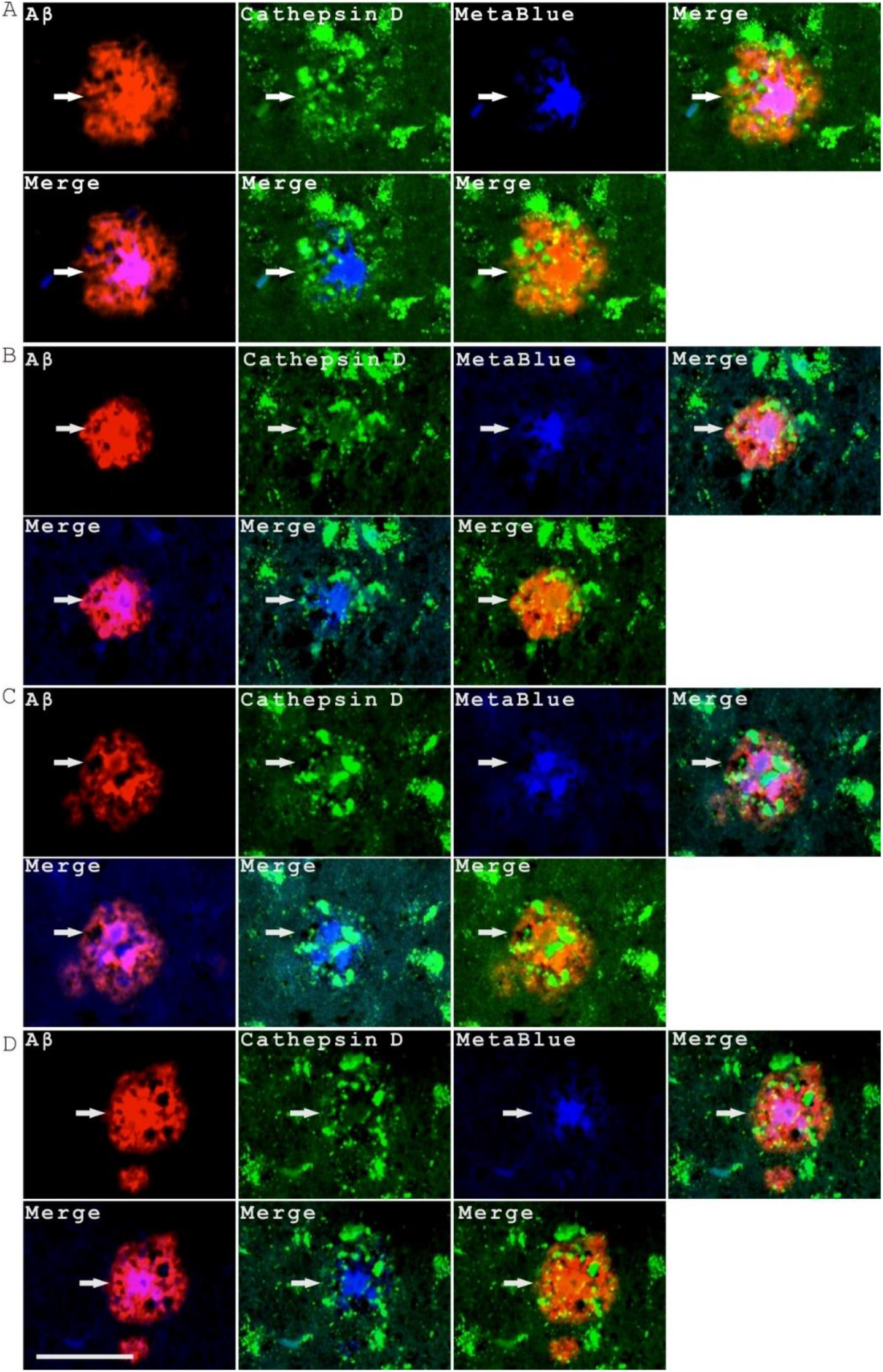
Cathepsin D, senile plaque Aβ staining and their relations to central plaque MetaBlue auto fluorescence. Four senile plaques (**A-D,** indicated with arrows) all showed that strong lysosomal Cathepsin D staining wrapped MetaBlue auto fluorescence staining in the center while senile plaque Aβ mostly co-localized with blue auto fluorescence staining. In addition, senile plaque Aβ co-localized with a weak diffusive Cathepsin D staining. Scale bar, 50 μm.

In this study, the authors used the “disintegration” and “disappearance” of nuclei staining as the most important marker of neural cell death since their Caspase-3 staining yielded negative results. Because the blue fluorescence in senile plaques might not come from nuclear signals at all, their interpretation of neuronal cell death needs to be re-examined. The author stated that “β-amyloid cored plaques, detected with β-amyloid antibodies, showed nearly 1:1 coincidence with a single PANTHOS neuron exhibiting a central nucleus.”, considering the dense-core plaques as individual abnormal neurons with degrading nucleus in the center. This statement probably is over-simplified and is not supported by other research data. First, according to our analysis, in human samples, the average diameter of typical senile plaques is more than 2 times of the size of average neuron cell bodies. The average area of senile plaques is around 5 times of the average area of neural cells. The average volume of a senile plaque is around 11 times of volume of an average neural cell body^2^. The chance of dense-core senile plaques being derived from individual neurons is low. Secondly, the centers of dense-core senile plaques contained blood vessels as previously reported^2,7,8^. Thirdly, the plaque cores contained Aβ and a large amount of blood-derived materials such as HBA, Hemin and ApoE^2^. In addition, from the evidence we showed above, Aβ-loaded plaque cores do not contain central cell nuclei but with central MetaBlue fluorescence materials instead. So called “PANTHOS” structures were not simple neuronal cell structures but rather complicated structures containing vascular and blood components. In our opinion, calling these plaques as “PANTHOS” neurons is not accurate and misleading.

## Conclusion

In summary, we think the nuclear staining data in this article should be verified and some interpretation of the data probably needs to be revised. We appreciate their great efforts in this interesting research about lysosomal defects in Alzheimer’s disease but the data and data interpretation should be more accurate especially when publishing in this high-profile journal of Nature Neuroscience. The material identity of MetaBlue is worthy of future investigation in order to elucidate the pathological mechanism of senile formation further.

## MATERIAL AND METHODS

### Tissue sections

AD patient frontal lobe brain paraffin tissue sections were provided by National Human Brain Bank for Development and Function, Chinese Academy of Medical Sciences and Peking Union Medical College, Beijing, China. This study was supported by the Institute of Basic Medical Sciences, Chinese Academy of Medical Sciences, Neuroscience Center, and the China Human Brain Banking Consortium. All procedures involving human subjects are done in accord with the ethical standards of the Committee on Human Experimentation in Shanghai Jiao Tong University and in Xinhua Hospital, and in accord with the Helsinki Declaration of 1975.

### List of antibodies for immunohistochemistry (IHC)

The following primary antibodies and dilutions have been used in this study: Aβ/AβPP (CST #2450, 1:200), Cathepsin D (Abcam ab75852, 1:200). The following secondary antibodies and dilutions have been used in this study: donkey anti-mouse Alexa-594 secondary antibody (Jackson ImmunoResearch 715-585-150, 1:400), donkey anti-rabbit Alexa-488 secondary antibody (Jackson ImmunoResearch 711-545-152, 1:400), and donkey anti-mouse Alexa-488 secondary antibody (Jackson ImmunoResearch 715-545-150, 1:400).

**Immunohistochemistry** was performed as described^9^. Briefly, paraffin sections were firstly deparaffinized by Xylene, 100% EtOH, 95% EtOH, 75% EtOH, 50% EtOH, and PBS washes. Sections were then treated with 10 mM pH6.0 sodium citrate or 10mM pH9.0 Tris-EDTA antigen retrieval solutions with microwave at first at high power for 5 minutes to get to the boiling point and then low power microwave for another 15 minutes. The sections were allowed to naturally cool down to room temperature. Then, the slides were blocked with TBST+3% BSA solution for 1 hour at room temperature. After blocking, the samples were incubated with primary antibodies at room temperature for 2 hrs followed by 5 washes of TBST. After that, the samples were incubated with fluorescent secondary antibodies overnight at 4 °C. The treated samples were washed again with TBST 5 times the second day and mounted with PBS+50% glycerol supplemented with PI (Sigma, P4170, 10 μg/ml) as a red nuclear fluorescence dye and ready for imaging. To analyze the blue auto fluorescence in those IHC experiments with also green and red fluorescent secondary antibodies, no nuclear fluorescent dye was included. IHC experiments without primary antibodies were used as negative controls. All experiments have been repeated twice taking from two different AD brain samples in order to verify the reproducibility of the results.

### Imaging analysis

The fluorescent images were taken with a CQ1 confocal fluorescent microscope (Yokogawa, Ishikawa, Japan). The blue autofluorescence of senile plaques (MetaBlue) can be imaged with the standard DAPI blue fluorescence channel using the CQ1 confocal microscope with the excitation wavelength of 405 nm, and the emission spectrum of 447 nm-460 nm.

## Acknowledgements

We want to thank the help from Dr. Ma Chao and Dr. Qiu Wenying for providing AD tissue sections from National Human Brain Bank for Development and Function, Chinese Academy of Medical Sciences and Peking Union Medical College, Beijing, China. Additionally, we want to thank Housheng Wang, Pingxin Liu, Xiaoyi Bao, and Jun Wang for excellent lab assistance.

## Author contributions

HF conceived the study, designed and supervised the experiments and wrote the manuscript; HF and JL performed the experiments; HF, JL, CZ, PD, GG, QG, XG and DC did the data analysis; All authors reviewed the manuscript.

## Data availability statement

The data that supports the findings of this study are available from the corresponding author upon request.

## References

1 Lee, J. H. et al. Faulty autolysosome acidification in Alzheimer’s disease mouse models induces autophagic build-up of Abeta in neurons, yielding senile plaques. Nat Neurosci 25, 688–701, doi:10.1038/s41593-022-01084-8 (2022).

2 Fu, H. et al. Senile plaques in Alzheimer’s disease arise from Abeta- and Cathepsin D-enriched mixtures leaking out during intravascular hemolysis and microaneurysm rupture. FEBS Lett, doi:10.1002/1873-3468.14549 (2022).

3 Dowson, J. H. A sensitive method for the demonstration of senile plaques in the dementing brain. Histopathology 5, 305–310, doi:10.1111/j.1365-2559.1981.tb01789.x (1981).

4 Diez, M., Koistinaho, J., Kahn, K., Games, D. & Hokfelt, T. Neuropeptides in hippocampus and cortex in transgenic mice overexpressing V717F beta-amyloid precursor protein--initial observations. Neuroscience 100, 259–286, doi:10.1016/s0306-4522(00)00261-x (2000).

5 Kwan, A. C., Duff, K., Gouras, G. K. & Webb, W. W. Optical visualization of Alzheimer’s pathology via multiphoton-excited intrinsic fluorescence and second harmonic generation. Opt Express 17, 3679–3689, doi:10.1364/oe.17.003679 (2009).

6 Gao, Y. et al. Imaging and Spectral Characteristics of Amyloid Plaque Autofluorescence in Brain Slices from the APP/PS1 Mouse Model of Alzheimer’s Disease. Neurosci Bull 35, 1126–1137, doi:10.1007/s12264-019-00393-6 (2019).

7 Kumar-Singh, S. et al. Dense-core senile plaques in the Flemish variant of Alzheimer’s disease are vasocentric. The American journal of pathology 161, 507–520, doi:10.1016/S0002-9440(10)64207-1 (2002).

8 Kumar-Singh, S. et al. Dense-core plaques in Tg2576 and PSAPP mouse models of Alzheimer’s disease are centered on vessel walls. The American journal of pathology 167, 527–543, doi:10.1016/S0002-9440(10)62995-1 (2005).

9 Fu, H. et al. Persisting and Increasing Neutrophil Infiltration Associates with Gastric Carcinogenesis and E-cadherin Downregulation. Sci Rep 6, 29762, doi:10.1038/srep29762 (2016).

